# Buzz Pollination: Investigations of Pollen Expulsion using the Discrete Element Method

**DOI:** 10.1101/2024.08.01.606085

**Authors:** Caelen Boucher-Bergstedt, Mark Jankauski, Erick Johnson

**Affiliations:** Department of Mechanical & Industrial Engineering, Montana State University, Bozeman, MT, USA

**Keywords:** buzz pollination, discrete element method (DEM), pollen, anther, vibrations

## Abstract

Buzz pollination involves the release of pollen from, primarily, poricidal anthers through vibrations generated by certain bee species. Despite previous experimental and numerical studies, the intricacies of pollen dynamics within vibrating anthers remain elusive due to the challenges in observing these small-scale, opaque systems. This research employs the discrete element method (DEM) to simulate the pollen expulsion process in vibrating anthers. By exploring various frequencies and displacement amplitudes, a correlation between the maximum jerk of anther walls and the initial rate of pollen expulsion is observed under translating oscillations. This study highlights that while increased vibration intensity enhances pollen release, the rate of increase diminishes at higher intensities. Our findings also reveal the significant role of pollen-pollen interactions, which account for upwards of one-third of the total collisions. Comparisons between poricidal and pseudoporicidal anther geometries suggest that pore size and shape also influence expulsion rates. This research provides a foundation for more comprehensive models that can incorporate additional factors such as cohesion, adhesion, and Coulomb forces, paving the way for deeper insights into the mechanics of buzz pollination and its variability across different anther types and vibration parameters.

## INTRODUCTION

Approximately 9% of all flowering plants have anthers with an apical pore, defined as a poricidal anther, that require buzzing bees to release pollen [6, 26]. The loss of animal pollinators, with bees accounting for the majority of flower visits, would reduce global food production by about 5-8% and necessitate a much larger increase in agricultural land to meet the demand with non-pollinated crops [1]. Moreover, around 75% of all crops would see a reduction in quality and yield without visiting pollinators [2, 19]. These changes would impoverish the nutritional balance of the human diet and disproportionately impact different parts of the world [1, 19]. As alternative pollination treatments are considered, *i*.*e*., hormonal, mechanical sonication, and robotic bees, a study of tomatoes (*Solanum lycopersicum L*.) showed a more than doubling of fruit weight when treated by bees that participate in buzz pollination rather than through mechanical methods [8].

The exploration of buzz pollination dates back to at least 1902, with laboratory investigations emerging in the 1970s [24]. After landing on a flower, bees bite the anther and rapidly contract and relax their flight muscles. This “buzzing” produces vibrations that translate and deform the anther causing pollen to be released [15, 22, 26]. Studies have revealed that bee-generated buzzes consist of a set of short bursts lasting around 0.5 seconds, with each individual burst varying from a few milliseconds to 0.1 seconds [26] and typically ranging in frequency between 100 to 400 Hz across various bee species [21, 26]. Exciting anthers with artificially generated vibrations, experiments have demonstrated a direct correlation between vibration amplitude, observed as anther displacement, and pollen release [21]. While frequency has been shown to have an impact on pollen release, its specific contribution has been difficult to decouple from the displacement amplitude and there may not be an *optimal* frequency for buzzing [21].

One of the largest hurdles in experimental work on buzz pollination is the difficulty to see the internal workings of flower anthers and pollen while vibrating due to the small scale, high speeds, and the opacity of the anther wall. Numerical models provide an alternative approach and efforts have been made to understand buzz pollination via statistical mechanics [7] and the billiards model [14], where single pollen particles reflect off of moving, rather than stationary, walls. As such, the focus of inquiry has allowed for a shift from observable, macro-scale phenomena, such as the amount of pollen expelled, towards finer-scale processes within the anther [7, 10, 21, 26].

Buchmann and Hurley [7] proposed that the rate of pollen expulsion would increase with an increase of stored, kinetic energy in the anther. To capture this process, their model splits the rate of energy change in the system into two parts. The first part theorized that as the anther walls vibrate back and forth there would be a net increase of energy in the system, from pollen-wall and pollen-pollen collisions. The second part suggested that as individual pollen are expelled they no longer contribute to the total energy, which causes the energy to decrease as they leave. The result of these considerations indicate that there is a maximum available energy that is proportional to the amount of pollen present, the anther geometry, and the excitation velocity, that decays over time with a non-linear decrease in pollen expulsion. Adding inelastic collisions would further slow the rate of pollen release.

Utilized by Hansen *et al*. [14], the billiards model offers additional insight into pollen release by directly calculating particle trajectory from particle-wall collisions. Controlling both the frequency(ies) and amplitude(s) of the anther motion allows for a more granular investigation into the energy applied to the system. This model neglected collisions between pollen particles by assuming the relative size of the particles is very small with respect to the anther volume and therefore the probability of interaction is likewise small. This assumption tacitly implies that increasing the number of particles in the anther has no effect on pollen expulsion. In addition, since the applied motion was side-to-side, the particles are seeded with an initial, random velocity to ensure they move along the length of the anther. Over the range of frequencies and anther displacements simulated, all but the lowest peak anther velocities saw 90% or more of the particles leave the anther in 0.5 s, demonstrating again a direct relationship between the movement of the system to pollen expulsion.

An alternative to the previous approaches, the discrete element method (DEM) has seen renewed interest due to the relatively easy access to high-performance computers and the ease at which the algorithm can be parallelized. DEM is able to simulate millions of particles simultaneously and the number of particles is only limited by the computational resources available. Similar to the billiards model, particle trajectories are solved through time. However, instead of being governed by reflections, the acceleration of DEM particles is calculated from the summed interactions with each other, wall boundaries, and additional external forces. DEM has been used to model granular solids to determine quantities too difficult to measure experimentally, such as the values and distributions of forces in spherical packing [11] and the self-assembly of spheres under one-dimensional vibrations [3]. DEM is a flexible tool that allows for a broader range of adjustable particle properties and conditions, such as non-uniform particle shapes and cohesive forces that can be broken, necessary for an expansive range of future work.

This paper leverages DEM to simulate and analyze the pollen-pollen and pollen-wall interactions in a vibrating anther, and subsequently, the rate of pollen expulsion. The results from a parameter sweep of frequencies and anther displacements demonstrate how DEM relaxes some of the assumptions that have been necessary to model pollen to date.

## THE DISCRETE ELEMENT METHOD

Unlike many traditional modeling approaches that rely on a fixed discretization, DEM does not require a mesh and instead centers the equations of motion on each particle [9]. The collection of surface and body forces acting on a particle of mass, *m*, are summed to find the total force, which then solves for that particle’s acceleration according to Newton’s second law (**F**_**tot**_ = *m***a**). The balance of rotational momentum is also considered in DEM, but by assuming pollen to be smooth, spherical grains [16] the rotational energy is vanishingly small in this problem and a detailed description is therefore ignored. With a small enough time-step, Δ*t*, the position, **x**, of every particle is updated by **x**_**new**_ = **x**_**old**_ + **a**Δ*t*^2^ and the process repeats over all particles until the simulation ends. Bold variables indicate vector quantities. The mesh-free DEM model in Siemens’ SimCenter STAR-CCM+ 2206 [23] was utilized for this study.

As the anther was translated side-to-side, the number of pollen-pollen and pollen-wall collisions and the pollen released from the anther were tabulated. Though not leveraged in this study, DEM also allows particle position-data to be collected over the entire duration of a simulation, which can be used to determine individual particle velocities and accelerations, and also aggregate data, such as the mean number of collisions before an individual pollen particle is released or the characteristic trends for pollen given a distribution of sizes.

Aerodynamic forces act on the surface of a particle and are caused through interactions with the surrounding fluid. These consist of the drag force from viscous shears and pressure distributions caused by moving in a fluid, the force from an additional pressure gradient, like buoyancy, and the added mass force, which is a contribution to the inertia of an accelerating, or decelerating, object as it displaces a fluid. In [7], aerodynamic forces had little to no impact on the amount of pollen expelled, but did reduce the distance traveled once outside the anther. Because the study only considers pollen inside the anther, we assume the pollen particles are traveling within a vacuum and, by extension, all surface forces cancel.

The body forces acting on the particles consist of the force from gravity, contact forces described by the chosen contact model, and the Coulomb force that accounts for electrostatic attraction or repulsion. A study by Bowker and Crenshaw [4] found an average positive charge across seven plant species of approximately 0.32 fC, yielding electrostatic forces that are of the same order as the gravitational force. The anther wall may shield pollen from electrostatic forces, but contactless transport has been observed externally to electrostatically charged butterflies and moths [13]. From preliminary simulations it was determined that gravitational acceleration had a negligible impact and mirrors the observations of [7] where the anther orientation did not significantly impact pollen expulsion. As such, gravity and electrostatic forces are also excluded in this initial study. Considering these simplifications, the total force acting on a pollen particle consists of only contact forces with other particles and walls.

### Contact Forces

The Hertz-Mindlin no-slip contact model is used to calculate the normal, *F*_*n*_, and tangential, *F*_*t*_, components of the reaction force from two colliding particles

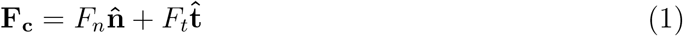

where the normal direction, 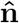, follows the line connecting their centers and the tangential direction, 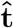, is parallel to a modified difference of the two particle velocities [12, 25]. FIG. 1 shows the contact force between two spheres, A and B. If multiple spheres interact at the same time, the force from each sphere-pair is summed into a cumulative contact force.

**FIG. 1.**
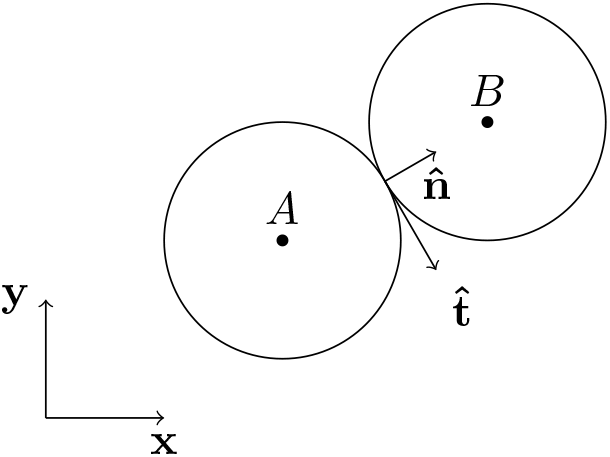
Simple collision of two particles with tangential and normal force components.

The normal and tangential force components are calculated using Equations 2a - 2c and 3a - 3c, respectively, with the forces combining the response from the material stiffness, *K*, and damping, *N*. Stiffness quantifies the force required to stretch or compress a particle by a unit length and damping refers to the force that opposes how quickly this occurs, scaled by the coefficient of restitution, *C*_*rest*_. This coefficient is a non-dimensional number that represents the fractional amount of kinetic energy that remains after a collision, with *C*_*rest*_ = 1 being perfectly elastic and all the energy being absorbed when *C*_*rest*_ = 0. This is seen as the material damping terms vanish when collisions are perfectly elastic. The tangential force is conditionally dependent on the static friction coefficient *C*_*fs*_.

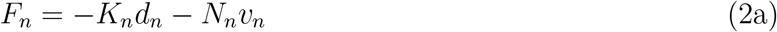

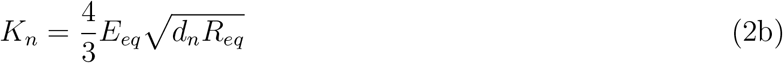

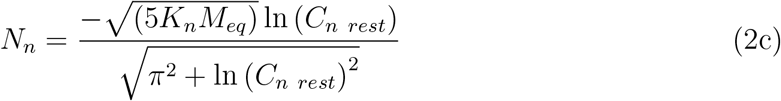

Because of the finite scale of the time-step the geometry of the two particles will intersect by a small amount, *d*, in each direction. The relative velocity between the two particles, ***v***, is split into normal and tangential components, with the latter deriving the tangential direction 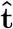. The subscripts *n* and *t* denote normal and tangential components, respectively, for any of the terms above.

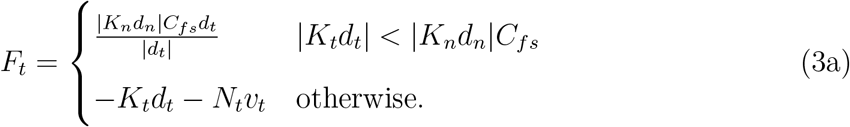

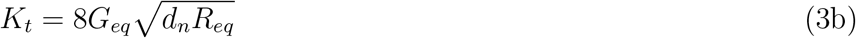

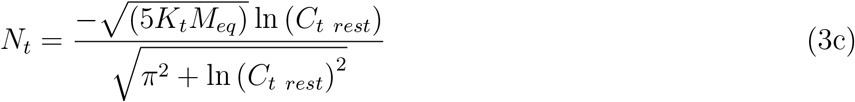

Equations 2 and 3 utilize equivalent physical values described by Equations 4b - 4d,

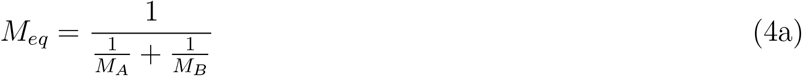

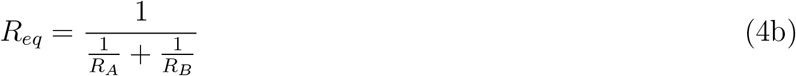

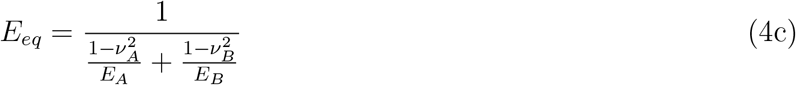

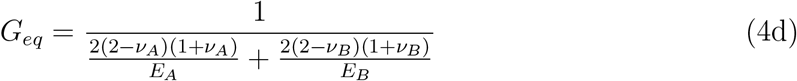

where spheres A and sphere B can have distinct mass *M*, radii *R*, and material properties: *E* is the Young’s modulus and *v* the Poisson’s ratio. The shear modulus, *G*, can be calculated from the others. When applying the Hertz-Mindlin no-slip contact model to a sphere and a wall, the wall radius and mass are assumed to be infinite *R*_*wall*_ = ∞ and *M*_*wall*_ = ∞.

### The Simulation Environment

Simplifying the anther to a box, two geometries were created: a poricidal design where a finite-sized pore is placed at the apical end, inspired by the paper by Buchmann and Hurley in 1978 [7], and a pseudoporicidal anther modeled as an extruded 2D shape, resulting in a slit at the apical end that matches the research done by Hansen *et al*. [14]. Both anther geometries were defined with the width and length being *a* = 0.69 mm and height *b* = 5.07 mm. Additionally, a hole at the top of the anther was dimensioned with *h* = 0.22 mm, such that the poricidal model was a square occupying 10.2% of the total top area and the slit of the pseudoproricidal model accounting for 31.9%. The anther geometry and pore shapes can be seen in FIG. 2.

**FIG. 2.**
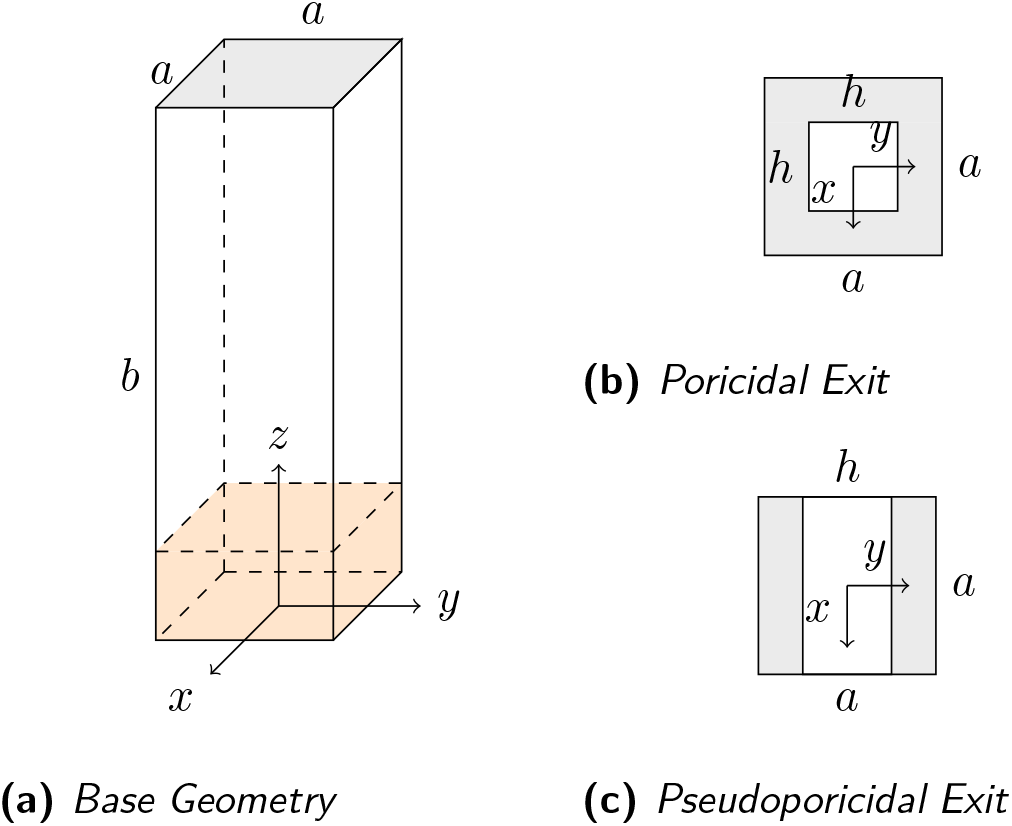
Simplified geometry representation. Shaded area represents region subject to particle injection.

The anther is subjected to a strictly translational motion along the y-axis of FIG. 2, in which the position of the anther wall is described by Equation 5,

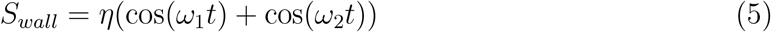

and subsequently the velocity of the anther wall is described by Equation 6,

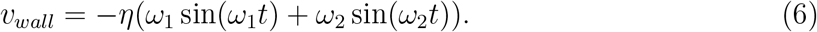

where *t* represents time, *η* is a displacement amplitude. Modeled after the beat frequency of [14], *ω*_1_ is the varied frequency and *ω*_2_ is a stationary frequency. The varied frequencies in the parameter ranged from 200 Hz to 1600 Hz in increments of 350 Hz, with the addition of 150 Hz to confirm a low-frequency trend. The peak displacement amplitudes, *i*.*e*., 2*η* due to the beat frequency, ranged from 0.4 mm to 2 mm in increments of 0.4 mm. A constant value of 390 Hz was used for *ω*_2_.

In each of the 60 simulations, 10,000 pollen particles were randomly seeded in the bottom 5% of the anther and represented by the shaded region in FIG. 2. The pollen particles started with a zero initial velocity and properties given in Table I. The Young’s modulus of pollen varies based on the hydration of the individual particle and the value selected is based on work done by [20]. Simulations ran for 0.5 s with a time-step of Δ*t* = 1 *µs*.

**TABLE I.** Pollen properties [5]^∗^ [18]^†^ [20]^‡^ and simulation values.

| Variable | Value |
| --- | --- |
| $E^{\ddagger}$ | 1.6 GPa |
| $\nu^*$ | 0.45 |
| $\rho$ | 1000 kg/m<sup>3</sup> |
| $R^{\dagger}$ | 0.01 mm |
| $C_{n \text{ rest}}$ | 1.0 |
| $C_{t \text{ rest}}$ | 1.0 |
| $v_0$ | 0 m/s |

## RESULTS AND DISCUSSION

Buzzing amplitude and frequency are known to be the principal drivers of pollen release, but the amount of pollen expelled as the lone measure has not drawn stronger conclusions. For all the conditions simulated, particle expulsion of 99% or more was achieved within 0.5 seconds. These results are in good agreement with those of [14], except at the lowest frequency and amplitude combinations where the billiards model sees the number of pollen released fall off rapidly and is closer to 40-50% at 400 Hz and 0.2 mm peak displacement. This smaller percentage is closer to the amount expelled (57.8%) in experiments of *Bombus impatiens* during their first visit to a tomato flower [17]. However, the billiards model ignores particle-particle interactions, which account for a significant number of the total collisions and anticipated to increase the amount of pollen released.

Starting from rest, particles are pushed by the wall in the direction of *S*_*wall*_ but quickly transition to collisions with each other. These particle-particle collisions rapidly diffuse the kinetic energy within the anther and introduce a velocity component that is no longer aligned with *S*_*wall*_, allowing particles to move towards the exit. After a short period of time, the distance between particles is sufficiently large that particle-wall interactions become dominant. As the net energy within the anther increases, pollen particles begin to escape the anther. Then, as a consequence of both the distance between particles and particles leaving through the exit, the number of new particle-particle collisions asymptotes to zero first. Finally, the particle-wall collisions continue until the anther empties. This pattern is observed across all simulations. The pseudoporicidal exit shows fewer total number of interactions and is likely due to the larger available exit area, which reduces the chance of a particle bouncing back towards the base of the anther. An example of the total number of particle-particle and particle-wall collisions is shown in FIG. 3 for *η* = 0.4 mm and *ω*_1_ = 150 Hz. Interestingly, particle-particle collisions accounted for an average of 30.03 *±* 1.32% and 39.50 *±* 2.67% of all collisions in poricidal and pseudoporicidal anthers, respectively, across the parameter sweep.

**FIG. 3.**
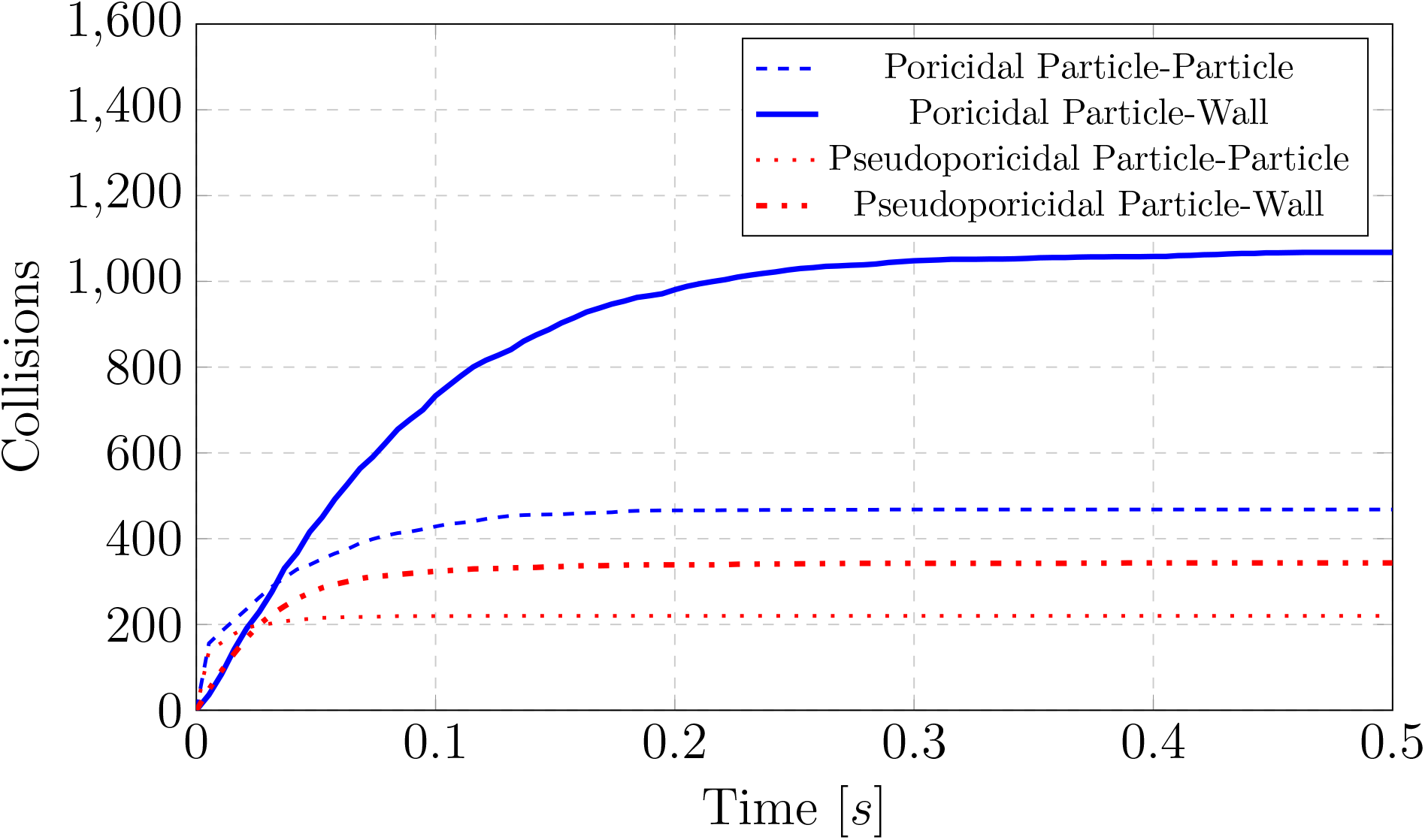
Comparison of cumulative particle interactions between poricidal and pseudoporicidal systems during the solution time of 0.5 seconds.

It is suspected that the difference in the amount of pollen expelled between the numerical models and experiments is principally a result of the neglected material damping. As derived in [7], the restitution coefficient plays a significant role in the available kinetic energy within the anther, where setting *C*_*rest*_ = 0.1 has a three orders-of-magnitude reduction in the normalized kinetic energy versus the perfectly elastic system. While values of *C*_*rest*_ can be calibrated to provide a better fit between models and the physical world, more experimental measurements of pollen and pollenkitt are needed. Additional losses, such as the breakup of pollen clusters and aerodynamic loads, would further reduce the number of pollen released over a given period of time.

The increase and decrease in kinetic energy is also captured in the total number of particles expelled from the anther over time, with a representative time-history shown in FIG. 4. For both the poricidal and pseudoporicidal geometries, particles are released faster when more particles are present and slower as fewer and fewer particles are present. Whether a result of the reduced kinetic energy available with a smaller number of particles or their distribution throughout the entire anther at later times, this slowing of particle release is observed through smaller dosings with repeated visits [17]. Though unsurprising, larger openings are also less restrictive and allow more particles to leave sooner.

**FIG. 4.**
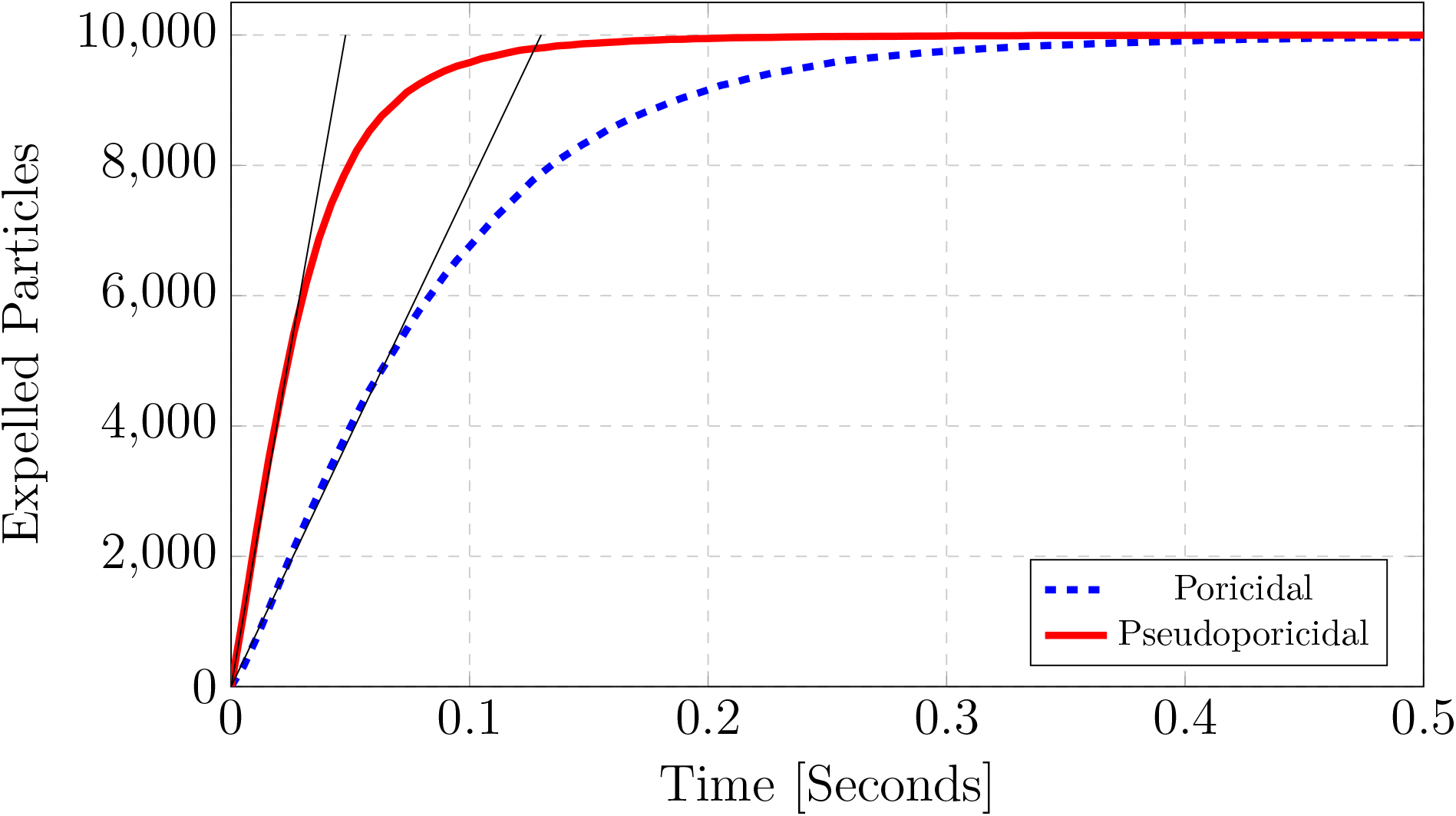
Examples of both poricidal and pseudoporicidal geometries pollen expulsion at η = 0.4 mm and ω_1_ = 150 Hz. The thin lines designate the initial expulsion rate of the particles,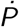, seen in FIG. 5.

While there was no difference in the amount of pollen released over the 0.5*s* of each simulation, a linear slope was observed for the first 50% of the total number of particles released. This initial pollen expulsion rate, 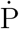 with units 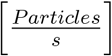 and seen as the thin, straight lines in FIG. 4, varied across the different parameter combinations, with the smallest rate corresponding with the smallest frequency and amplitude pair and the largest rates with the largest parameter combinations.

In discussing anther dynamics, only displacement amplitude and frequency are independent variables, *i*.*e*., velocity (Equation 6) and acceleration are time derivatives of the displacement profile and intrinsically coupled to both *η* and *ω*. Rather than considering these variables at odds with each other, all of the time derivatives of displacement collapse into a single variable. Here we introduce the maximum jerk, 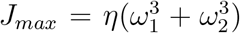, to describe how vigorously the anther shakes, or the change in acceleration over time. The initial rate of pollen expulsion is plotted against the maximum jerk and shown in FIG. 5 for all simulations. These results show a direct increase in pollen expulsion as *J*_*max*_ increases, which slows as *J*_*max*_ reaches significantly large values. This suggests that it does not matter whether frequency or amplitude drive this increase and that there are diminishing returns to how quickly pollen is released as more power is exerted to shake the anther. Moreover, five additional simulations were run with a single, continuous frequency with the poricidal geometry and the results fall right on top of these curves (shown as black Xs in FIG. 5). The implication is that, for purely translational motions, the specifics of how an anther is shaken is less important than how aggressively.

**FIG. 5.**
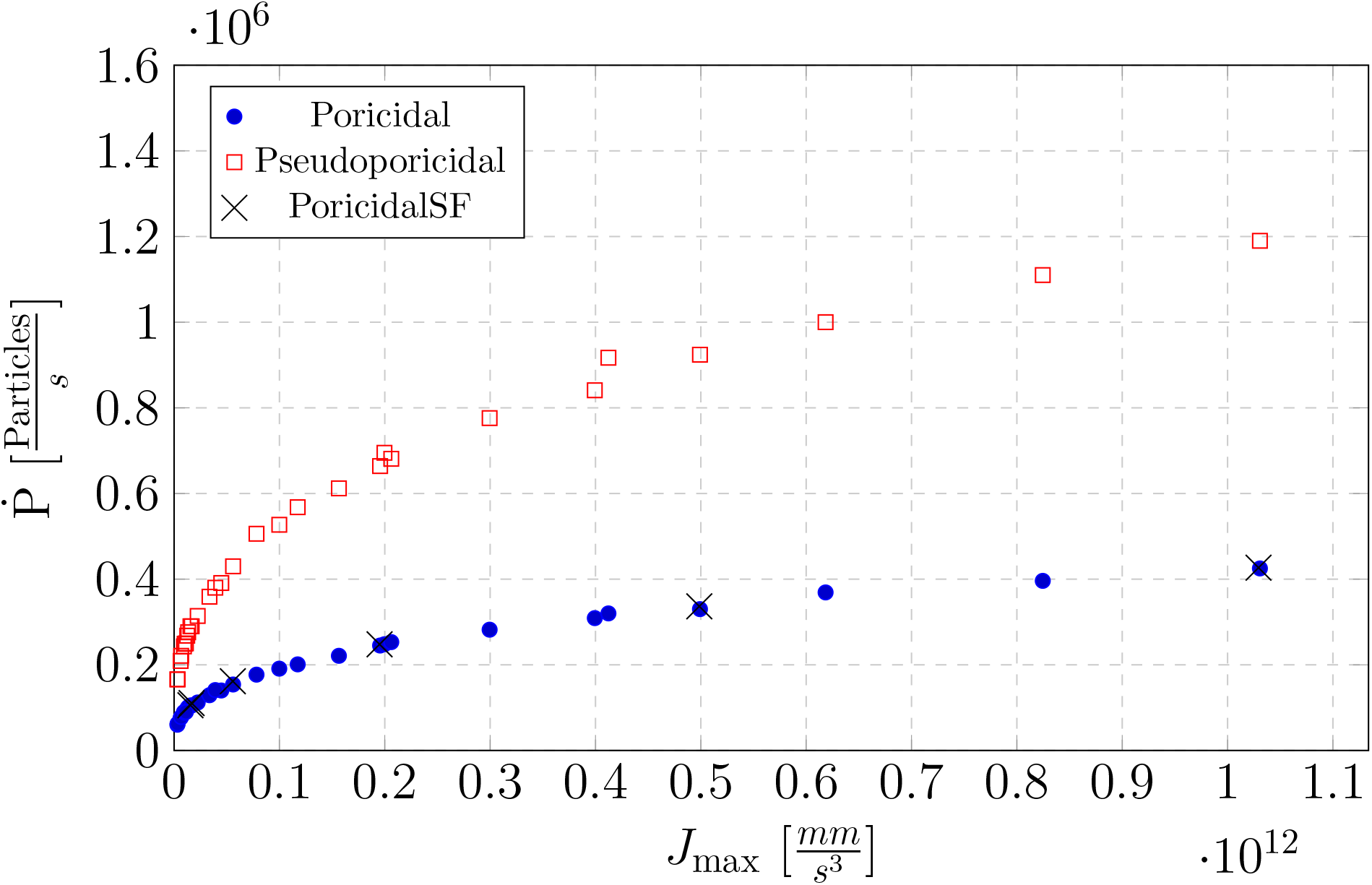
The initial pollen expulsion rate for both poricidal and pseudoporicidal anthers as a function of the maximum jerk of the anther walls. PoricidalSF indicates points where a different vibration profile, S_wall_ = η cos(ω_1_t), was utilized to test the effect of a single frequency.

## CONCLUSIONS

This research has utilized the discrete element method to begin exploring the complex dynamics of pollen within flower anthers during buzz pollination, subjected to simple trans-lating oscillations. Our findings reveal a strong correlation between the maximum jerk, *J*_*max*_, of anther walls and the initial rate of pollen expulsion. The simulations indicate that while increased rates of expulsion are achievable across a spectrum of values, further increases in vibration intensity show diminishing returns. This relationship suggests an “optimal” vibration profile may not exist and that non-translating deformations need to be investigated. The simulations also show that by including particle-particle interactions, we are able to expel pollen starting at rest using a purely transverse motion. Furthermore, approximately one-third of collisions that occurred within the anthers were between particles, contradicting previous estimates that particle-particle collisions were unimportant. Differences between the poricidal and pseudoporicidal anther exits indicate a dependence on the pore size and shape when quantifying the pollen expulsion rate.

This method of modeling also leaves room for including currently omitted factors, such as cohesion/adhesion and Coulomb forces, with only minor changes to the current approach. Further investigations could extend the range of modeled anther and pollen types and refine predictive models based on these findings. Additionally, expanding the variety of anther geometries examined could provide deeper insights into the mechanical interactions within buzz pollination.

## ACKNOWLEDGEMENTS

This material is based upon work supported by the National Science Foundation (NSF) (Grant No. CMMI-2221908). Any opinions, findings, and conclusions or recommendations expressed in this material are those of the authors and do not necessarily reflect the views of the NSF.

